# Gigantic Animal Cells Suggest Organellar Scaling Mechanisms Across a 50-fold Range in Cell Volume

**DOI:** 10.1101/2023.08.30.555588

**Authors:** Alexander Nichols Adams, Bradford Julian Smith, Thomas John Raad, Rachel Lockridge Mueller

**Affiliations:** Department of Biology, Colorado State University, Fort Collins CO 80523-1878; Department of Bioengineering, College of Engineering, Design & Computing, University of Colorado Denver | Anschutz Medical Campus, Aurora, CO, United States; Section of Pulmonary and Sleep Medicine, Department of Pediatrics, University of Colorado Denver, Anschutz Medical Campus, Aurora, CO 80045, USA

**Keywords:** Organelle Size, Salamanders, Genome Size, Cell Size, Scaling Relationships, Evolutionary Cell Biology

## Abstract

Across the tree of life, cell size varies by orders of magnitude, and organelles scale to maintain cell function. Depending on their shape, organelles can scale by increasing volume, length, or number. Scaling may also reflect demands placed on organelles by increased cell size. The 8,653 species of amphibians exhibit diverse cell sizes, providing a powerful system to investigate organellar scaling. Using transmission electron microscopy and stereology, we analyzed three frog and salamander species whose enterocyte cell volumes range from 228 to 10,593 μm^3^. We show that the nucleus increases in radius while the mitochondria increase in total network length; the endoplasmic reticulum and Golgi apparatus, with their complex shapes, are intermediate. Notably, all four organelles increase in volume proportionate to cell volume. This pattern suggests that protein concentrations are the same across amphibian species that differ 50-fold in cell size, and that organellar building blocks are incorporated into more or larger organelles following the same “rules” across cell sizes, despite variation in metabolic and transport demands. This conclusion contradicts results from experimental cell size increases, which produce severe proteome dilution. We hypothesize that salamanders have evolved the biosynthetic capacity to maintain a functional proteome despite a huge cell volume.

## Introduction

Cell size shows spectacular diversity across the Tree of Life, with unicellular organisms – where cell size is body size – varying up to a 1,000,000-fold in volume (Mueller 2015; Smith 2017; Malerba and Marshall 2021). In multicellular organisms – where cell size shapes the basic building blocks of tissues and organs – cells show size variation of around 1,000-fold (Gregory 2001, 2023). As cell size changes, assuming shape remains constant, the surface area to volume ratio (SA:V) scales in predictable ways dictated by the cell’s shape. Previous studies of small cells have shown how this scaling of SA:V impacts many aspects of a cell’s biology including movement across the cell membrane, metabolic flux, the reach of microtubules, and nutrient exchange (Marshall et al. 2012; Marshall 2020).

As cell size increases, organelles must scale to maintain overall cell functionality (Reber and Goehring 2015). Within the natural range of cell size, scaling among smaller cells has been shown to happen in different ways: by increasing in volume, increasing of linear network length, or increasing in copy number (Chan and Marshall 2010). Specific examples include organelles like the nucleus and nucleolus increasing in volume (Noel et al. 1971; Jorgensen et al. 2007), while mitotic spindles and mitochondria increase in network length (Hara and Kimura 2009; Rafelski et al. 2012) and multiple-unit organelles like peroxisomes increase in copy number (Titorenko et al. 2000). Other organelles of more complex shape like the endoplasmic reticulum (ER) and Golgi apparatus have been the focus of less research, and their scaling patterns are less clear (Chan and Marshall 2010; Marshall 2015).

As with cells, as organellar volume increases, the surface area scales differently depending on shape; an organelle shaped like a sphere shows a SA:V ratio that scales as r^2^ : r^3^, where r is the radius of the sphere. In contrast, an organelle shaped like a tube maintains a SA:V ratio of 1 : 1 as it grows in length (Chan and Marshall 2010). These two basic structures account for most organelles and reflect their functions; the nucleus is a large space that houses DNA and its associated molecules, and it tends to be more spherical. The mitochondria weave throughout the cell providing energy, and they tend to be more tubular (Chan and Marshall 2010; Marshall 2015). The ER and Golgi apparatus, however, are made up of cisternae, which are flattened membrane vacuoles that share attributes of both structures (Day et al. 2013; Schwarz and Blower 2016).

In plant species and some fungal species, cell size can be heavily impacted by large fluid-filled vacuoles, which can comprise 30% to 90% of the cell volume; additionally, storage plastids also contribute to intracellular volume (Klionsky et al. 1990; O’Connor et al. 2010). The size of these vacuoles can be highly variable and can cause overall cell size to shift dramatically (Chan and Marshall 2014; Tan et al. 2019); although the cells are enclosed by a cell wall, the walls themselves have the ability to expand and contract rapidly to accommodate changes in volume (Cosgrove 1993; Marshall et al. 2012).

In contrast, animal cells do not contain these storage organelles and plastids (Wise 2007; O’Connor et al. 2010), meaning their cell volume is almost exclusively derived from cytosol and membrane-bound organelles used for intra- and intercellular functions other than storage. In addition, animal cells closely regulate their osmotic gradient and maintain tight control on their cell size (Alberts et al. 2017; Cooper and Adams 2022). Thus, animal cells do not undergo rapid size change outside of growth and size reduction associated with the cell cycle, remaining at a more stable equilibrium size than plant cells. Two animal groups are composed of somatic cells at the farthest end of this size spectrum — salamanders and lungfishes. Salamanders are one of the three extant clades of amphibians, which include 8,653 species (Amphibiaweb 2023) with genome sizes ranging from 0.95 Gb −120 Gb; because genome size and cell size are correlated, the clade also exhibits a huge range of cell sizes (Gregory 2023).

Amphibians have been used for decades as a model taxon to examine how cell size affects an organism at the tissue and organismal levels (Hanken 1983; Roth et al. 1994; Womack et al. 2019; Decena-Segarra et al. 2020; Miller et al. 2020; Itgen et al. 2022). We continue to leverage this natural diversity of cell size across extant amphibians to quantify how organelles scale across a ~50-fold difference in cell volume using transmission electron microscopy (TEM) and stereology (Howard and Reed 2004; Russ and Dehoff 2012; Winey et al. 2014). We focus our analyses on the nucleus, mitochondria, endoplasmic reticulum (ER), and Golgi apparatus. The nucleus is the primary site for DNA and RNA synthesis, as well as the cell’s central hub for detecting deformations in cell shape (Venturini et al. 2020). Mitochondria are a network of membrane-bound organelles whose functions include ATP synthesis, apoptosis, and signaling (Brand et al. 2013). The ER is a network of membranous sacs and tubules that functions in lipid synthesis and protein synthesis, modification, and sorting (Schwarz and Blower 2016). The Golgi apparatus is a collection of stacked membrane-bound organelles that functions in protein sorting and trafficking and carbohydrate and lipid synthesis (Day et al. 2013). Larger cell size increases intracellular trafficking distance, as well as increasing per-cell demand for ATP, transcript and protein production, and lipid and carbohydrate synthesis (Guo and Fang 2014); thus, functional demands on all of these organelles are likely impacted by evolutionary increases in cell size. Taken together, our results reveal how organelles scale to maintain cell functionality at the extremes of animal cell size.

## Materials and Methods

### Tissue Sampling, Fixation, Staining, and Imaging

Intestinal tissue was chosen for analysis as it is made up of only four cell types, and 80 percent of the total cell population is enterocytes, resulting in a relatively homogenous population of cells (De Santa Barbara et al. 2003). Three species of amphibians were chosen that span much of the range of amphibian genome and cell sizes: the western clawed frog *Silurana tropicalis* (genome size = 1.2 Gb), the northern gray-cheeked salamander *Plethodon montanus* (genome size = 35 Gb), and the western waterdog *Necturus beyeri* (genome size ~100 Gb based on congeners that range from 80.5-120.6 Gb). *Silurana tropicalis* were obtained from a lab-reared colony following standard husbandry conditions and *Necturus beyeri* were obtained commercially. *Plethodon montanus* were field collected between May and August of 2018 in Avery County, North Carolina under the wildlife collection license # 18-SC01250 issued by the North Carolina Wildlife Resources Commission. One individual was sampled per species, and all specimens were euthanized in MS222. Work was carried out in accordance with Colorado State University (*P. montanus, N. beyeri*) and University of Wyoming (*S. tropicalis*) IACUC protocols (17-7189A and 20200714DL00443-01, respectively).

Intestinal tissue from each individual was dissected and immersion fixed in 2.5% glutaraldehyde/2% formaldehyde. The tissues then underwent secondary fixation and staining in 1% OsO_4_ in a 0.1 M cacodylate buffered solution followed by embedding in PELCO Eponate 12 epoxy (Cushing et al. 2014). Thin sections (60-80 nm) of resin-embedded samples were cut using a Leica UCT ultramicrotome, collected onto Formvar-coated TEM slot grids, and poststained with 2% aqueous uranyl acetate followed by Reynold’s lead citrate.

Sample preparation, fixation, and mounting were done at Colorado State University. The samples were then sectioned, stained, and imaged at the University of Colorado, Boulder Electron Microscopy Services Core Facility. Sections were imaged using a Tecnai T12 Spirit transmission electron microscope, operating at 100 kV, with an AMT CCD digital camera. *Silurana tropicalis* and *P. montanus* tissues were imaged at 9,300x direct magnification. *Necturus beyeri* tissue was imaged at 6,800x direct magnification because the larger cell sizes could not be captured at the higher magnification. All images were evaluated for quality, ensuring intact tissues undamaged by the fixation process.

### Stereological Approach

Stereology uses 2-dimensional image sampling protocols that allow the estimation of surface area and volume of 3-dimensional shapes through unbiased sampling using grids (Howard and Reed 2004; Russ and Dehoff 2012). Grids are superimposed randomly onto preselected, non-overlapping TEM images. Depending on the type of probes used in the grid and how the user measures the probes’ interactions with the images, the volume, surface area, length, and number of 3-dimensional structures can be estimated from the 2-D TEM images.

For the individual representing each species, 60 TEM images were randomly selected, and each image was overlayed with a grid of probes using the IMOD image processing program (Kremer et al. 1996). Volume fraction of organelles per cell (point probes), as well as surface area density (line probes), were measured for nucleus, mitochondria, ER, and Golgi apparatus using the 3dmod stereology plugin in IMOD (Noske 2010).

### Organelle Volume Fraction Estimation

For measuring volume fractions of the organelles (i.e. the volume of each type of organelle per total cell volume), each of the 60 images for each individual was fitted with a 7 × 7 grid using crosshair probes (points) for a total of 2,940 probes per species. Next, each crosshair was visually identified and manually assigned to one of the following categories: nucleus, mitochondria, ER, Golgi apparatus, or cytoplasm (which included cytosol and other non-focal organelles). If a crosshair fell on a part of the image that included damaged cellular material or extracellular material, it was categorized as “not in bounds” and was removed from the data set. The center of the crosshair was used to define the object category, and only one class was permitted per point (Howard and Reed 2004; Russ and Dehoff 2012). A denser grid was used for the Golgi apparatus due to the rarity of the organelle; the same 60 images per individual were overlaid with 18 × 18 grids for a total of 19,440 crosshair probes.

### Organelle and Cell Surface Area Density Estimation

For measuring organelle and cell surface area densities (i.e. surface area per unit of cell volume), the same 60 images per individual were used as for volume fraction estimates. The nucleus, mitochondria, ER, and plasma membrane were measured simultaneously using alternating cycloid probes in a 2 × 2 grid per image for a total of 240 probes. The Golgi apparatus was again measured using a denser grid due to the organelle’s scarcity, using a 6 × 6 grid of alternating cycloids on each image for a total of 2,160 cycloids per species. The cycloids were visually identified and manually assigned as being “in bounds” and “not in bounds” as in the volume fraction estimation. From there, cycloids were marked with intercepts each time a cycloid interacted with the boundary of an organelle or cell; if a cycloid went entirely through an organelle, then an intercept was marked both entering and leaving the structure. Each intercept was categorized as one of the four groups of organelles from the volume fraction estimation or as the cell boundary; cytoplasm was not used in surface area density estimation. Cycloids that were fully inside an organelle and did not contact the outside of the organelle were not marked with intercepts. The surface area density estimates were then calculated from these results with the equation 2*Intercepts/Total Length of all “In Bound” Cycloids (Howard and Reed 2004; Russ and Dehoff 2012).

### Organelle and Cell Surface Area to Volume Ratio Estimation

Organelle and whole cell SA:V ratios for each species were calculated by dividing the estimates for surface area density by the volume fraction estimates for each image. Because organellar structures are so variable in shape (e.g. the nucleus and mitochondria exhibiting spherical vs. tubular network shapes) (McCarron et al. 2013; Malerba and Marshall 2021), their SA:V ratios scale in different ways with increases in organellar size (Chan and Marshall 2010; Marshall 2020). Thus, SA:V estimates across species act as a proxy for organelle shape.

### Organelle and Cell Absolute Volume Estimation

Nuclear volume was estimated using the Nucleator probe (Gundersen et al. 1988) in the Visiopharm VIS stereology software (version 2017.7). The Nucleator randomly assigns two perpendicular rays that radiate outward from a fixed point in the nucleus, defined to be the nucleolus, and uses these rays and their intersection points with the nuclear membrane to estimate mean nuclear volume. 100 nuclei were analyzed from each species. Then, using the proportional estimates of nuclear volume obtained from IMOD (Kremer et al. 1996), cell volume and surface area, and all other organelle volumes and surface areas, were extrapolated from nuclear volume.

### Among-Species Differences in Organelle Volume Fraction, Surface Area Density, and Surface Area to Volume Ratio

We tested for associations between cell size (i.e. species) and 1) organelle volume fraction, 2) organelle surface area density, and 3) organelle surface area to volume ratio using ANOVA, followed by a pairwise Tukey post-hoc method of comparison. We carried out all analyses in RStudio (Rstudio Team 2021) using R packages emmeans (Lenth 2021), car (Fox and Weisberg 2011) and lme4 (Bates et al. 2015).

## Results

### Organelle Volume Fraction

Across the three species, the cytoplasm made up the largest fraction of cell volume (0.48 – 0.57), followed by the nucleus (0.27 – 0.36), mitochondria (0.09 – 0.12), ER (0.04 – 0.07), and Golgi apparatus (0.002 – 0.003) (Fig. 1). None of the four organelles showed any statistically significant differences in volume fraction across species based on the ANOVA (p > 0.05). The cytoplasm volume fraction was statistically significantly different between *S. tropicalis* and *P. montanus* (ANOVA p = 0.002; Tukey p = 0.002). All mean volume fractions and standard errors are listed in Table 1. All p-values are listed in Table 2.

**Figure 1:**
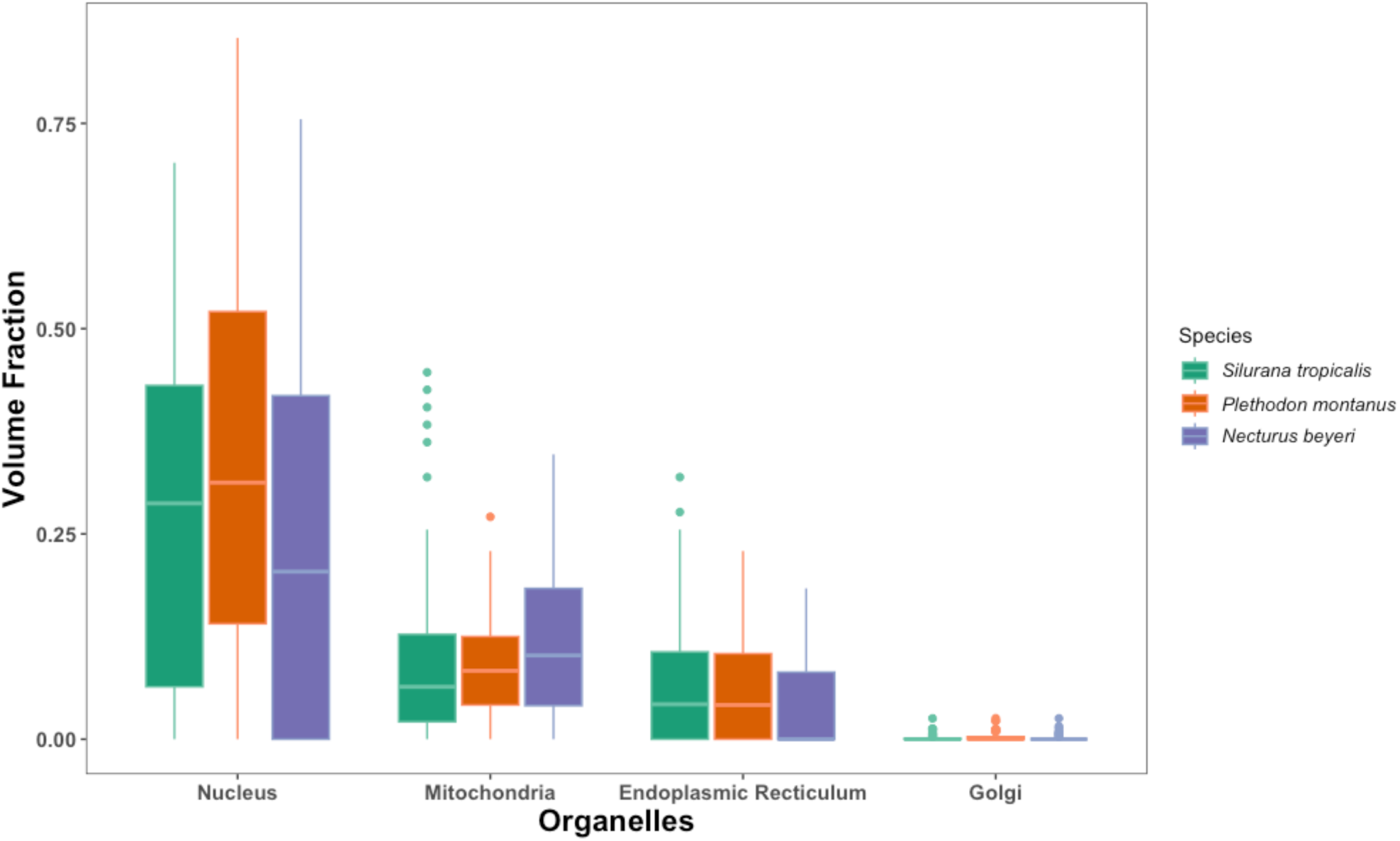
Volume fraction of each organelle. Species are shown left-to-right from smallest to largest cell sizes.

**Table 1:** Measurements of the cell and organelles (* Indicates significant difference p < 0.05)

| Measurement | Component | *Silurana tropicalis*<br>Mean (SE) | *Plethodon montanus*<br>Mean (SE) | *Necturus beyeri*<br>Mean (SE) |
| --- | --- | --- | --- | --- |
| Volume Fraction | Cytoplasm | **0.57\*** (0.02) | **0.49\*** (0.02) | 0.56 (0.02) |
|  | Nucleus | 0.27 (0.03) | 0.36 (0.03) | 0.27 (0.03) |
|  | Mitochondria | 0.10 (0.01) | 0.09 (0.01) | 0.12 (0.01) |
|  | Endoplasmic Reticulum | 0.06 (0.008) | 0.07 (0.008) | 0.04 (0.008) |
|  | Golgi Apparatus | 0.002 (0.0001) | 0.003 (0.0001) | 0.002 (0.0001) |
| Surface Area Density<br>($\mu\text{m}^2/\mu\text{m}^3$) | Cell | **0.93\*** (0.06) | **0.61\*** (0.06) | **0.27\*** (0.06) |
|  | Nucleus | **0.46\*** (0.04) | **0.25\*** (0.04) | **0.10\*** (0.04) |
|  | Mitochondria | 1.19 (0.05) | 1.01 (0.05) | 1.15 (0.05) |
|  | Endoplasmic Reticulum | **0.71\*** (0.08) | 0.57 (0.08) | **0.34\*** (0.08) |
|  | Golgi Apparatus | 0.07 (0.02) | 0.07 (0.02) | 0.03 (0.02) |
| Surface Area to Volume Ratio<br>($\mu\text{m}^2/\mu\text{m}^3$) | Cell | **0.97\*** (0.07) | **0.68\*** (0.07) | **0.31\*** (0.07) |
|  | Nucleus | **1.52\*** (0.20) | **0.83\*** (0.19) | **0.26\*** (0.19) |
|  | Mitochondria | 12.31 (1.8) | 11.54 (1.8) | 12.57 (1.8) |
|  | Endoplasmic Reticulum | 10.05 (1.9) | 7.50 (1.9) | 4.83 (1.9) |

**Table 2:** ANOVA p-values for volume fraction, surface area density, and SA:V ratio for the cell, cytoplasm, and organelles across species. All Tukey post-hoc p-values for between-species comparisons. (* Indicates significant difference)

|  | Across All Species<br>(p-value) | *S. tropicalis* –<br>*P. montanus*<br>(p-value) | *P. montanus* –<br>*N. beyeri*<br>(p-value) | *S. tropicalis* –<br>*N. beyeri*<br>(p-value) |
| --- | --- | --- | --- | --- |
| **Cytoplasm Volume Fraction** | **0.002\*** | **0.002\*** | 0.052 | 0.478 |
| **Cell SA Density** | **8.23E-11\*** | **0.002\*** | **0.001\*** | **<0.001\*** |
| **Cell SA:V Ratio** | **1.04E-08\*** | **0.018\*** | **0.001\*** | **<0.001\*** |
| **Nucleus Volume Fraction** | 0.102 | 0.233 | 0.110 | 0.925 |
| **Nucleus SA Density** | **5.14E-10\*** | **0.0002\*** | **0.014\*** | **<0.001\*** |
| **Nucleus SA:V Ratio** | **4.34E-05\*** | **0.034\*** | 0.093 | **<0.001\*** |
| **Mitochondria Volume Fraction** | 0.218 | 0.656 | 0.188 | 0.660 |
| **Mitochondria SA Density** | 0.585 | 0.606 | 0.688 | 0.990 |
| **Mitochondria SA:V Ratio** | 0.899 | 0.921 | 0.911 | 0.999 |
| **ER Volume Fraction** | 0.078 | 0.999 | 0.129 | 0.118 |
| **ER SA Density** | **0.007\*** | 0.446 | 0.124 | **0.005\*** |
| **ER SA:V Ratio** | 0.568 | 0.580 | 0.113 | 0.136 |
| **Golgi Volume Fraction** | 0.481 | 0.739 | 0.454 | 0.893 |
| **Golgi SA Density** | 0.320 | 1.000 | 0.388 | 0.396 |

### Surface Area Density of Cell and Organelles

All mean surface area densities and standard errors are listed in Table 1. Across the three species, the largest organellar surface area density is seen in the mitochondria (1.0 – 1.2 μm^2^/μm^3^), followed by the ER (0.3 – 0.7 μm^2^/μm^3^), nucleus (0.1 – 0.5 μm^2^/μm^3^), and Golgi apparatus (0.03 – 0.07 μm^2^/μm^3^) (Fig. 2). All 3 species show statistically significant differences in nuclear surface area density (p < 0.02; Table 2), with nuclear surface area density decreasing as cell size increases. The ER also shows a decrease in surface area density as cell size increases, with a statistically significant difference between *S. tropicalis* and *N. beyeri*, the smallest and largest cells (p < 0.007; Table 2).

**Figure 2:**
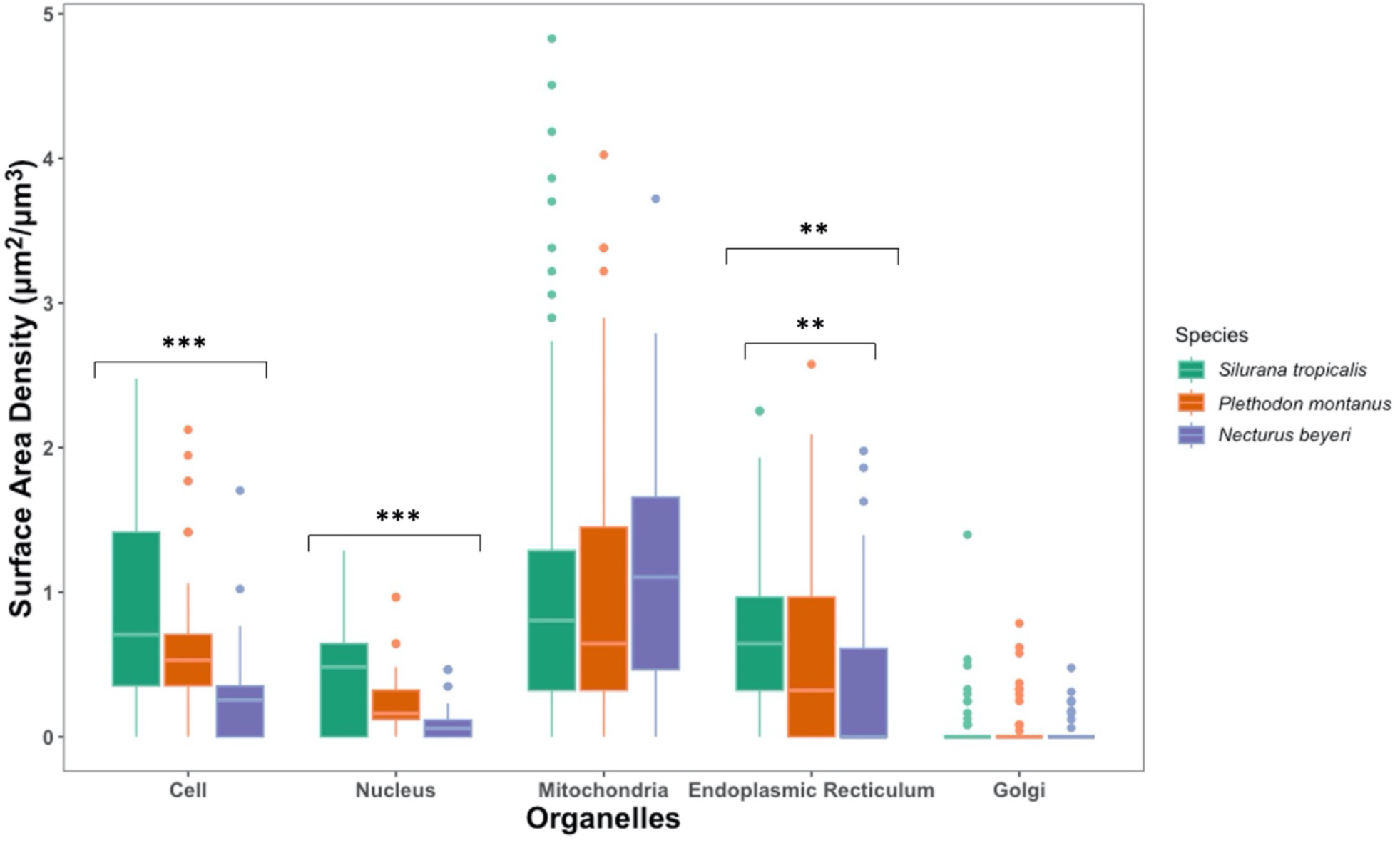
Surface area density for each organelle and the overall cell. Species are shown left-to-right from smallest to largest cell sizes. Groups with significant differences are marked with asterisks; the one pairwise comparison of the ER that shows a significant difference in the post-hoc test is indicated further (** p < 0.01, *** p < 0.001).

### Surface Area to Volume Ratio of the Cell and Organelles

All mean SA:V ratios for the cell and organelles and standard errors are listed in Table 1. Across the three species, SA:V ratio of the cell is statistically significantly different (p < 0.02; Table 2), decreasing from 0.97 μm^2^/μm^3^ in *S. tropicalis* to 0.31 μm^2^/μm^3^ in *N. beyeri*. As expected, the SA:V ratio of the nucleus also decreases from 1.52 μm^2^/μm^3^ to 0.26 μm^2^/μm^3^ as cell size increases and is statistically significantly different between *S. tropicalis* and *P. montanus* as well as between *S. tropicalis* and *N. beyeri* (p < 0.034; Table 2). In contrast, the mitochondria show no differences in SA:V ratio across species, consistent with the expectation of increasing the length of a tube-like network as cell size increases. The ER also shows no differences in SA:V ratio across species. Because of its scarcity, the Golgi was absent from many grids, so per-grid SA:V ratios were not calculated.

**Figure 3:**
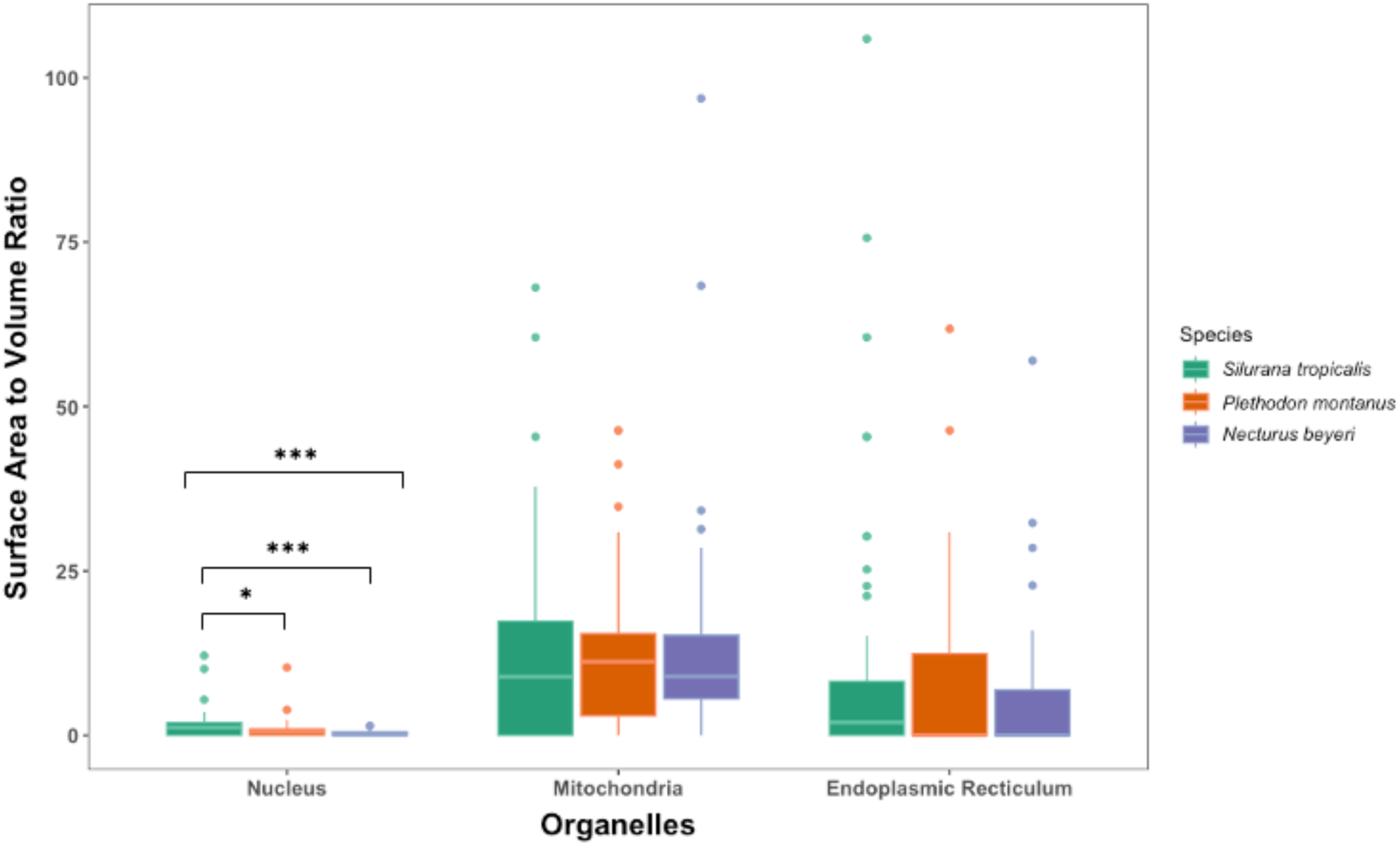
Surface Area to Volume ratio of the organelles. Species are shown left-to-right from smallest to largest cell sizes. Groups with significant differences are marked with asterisks, subgroupings that show significant differences in post-hoc tests are indicated further (* p < 0.05, *** p < 0.001)

### Estimates of Absolute Cell and Organellar Volume and Surface Area

All estimates of absolute volume and surface area for cells and organelles are listed in Table 3. Cell volume in *N. beyeri* is 46 times larger than *S. tropicalis.* The majority of cell volume estimates from diverse taxa in the literature are for erythrocytes; comparison of our enterocyte volumes with erythrocyte volumes for the same or congeneric species reveals that enterocytes are larger (*S. tropicalis* 228 μm^3^ vs. 122 μm^3^; *P. montanus* 3,191 μm^3^ vs 1,476 – 3,204 μm^3^ for congeners *P. cinereus* (20 Gb) and *P. dunni* (47.5Gb) (Gregory 2023)), but that our estimates are in keeping with other empirical estimates.

**Table 3:** Estimated volumes and surface areas extrapolated from Nucleator estimates of nuclear volume and point count estimates of volume fractions and surface area densities.

|  | *Silurana tropicalis* |  | *Plethodon montanus* |  | *Necturus beyeri* |  |
| --- | --- | --- | --- | --- | --- | --- |
| | Volume ($\mu\text{m}^3$) | Surface Area ($\mu\text{m}^2$) | Volume ($\mu\text{m}^3$) | Surface Area ($\mu\text{m}^2$) | Volume ($\mu\text{m}^3$) | Surface Area ($\mu\text{m}^2$) |
| Entire Cell | 228.2 | 212.3 | 3,190.9 | 1926.7 | 10,592.8 | 2,777.6 |
| Cytoplasm | 128.6 | N/A | 1,525.6 | N/A | 5,983.9 | N/A |
| Nucleus | 62.2 | 105.0 | 1,154.4 | 797.7 | 2,843.1 | 1,059.3 |
| Mitochondria | 22.6 | 271.6 | 292.6 | 3,222.8 | 1,301.9 | 12,181.7 |
| Endoplasmic Reticulum | 14.3 | 162.0 | 210.6 | 1,818.8 | 448.1 | 3,601.5 |
| Golgi Apparatus | 0.52 | 16.0 | 7.9 | 223.4 | 15.9 | 317.8 |

## Discussion

Marshall (2020) laid out five mechanistic models to explain the scaling of organelles with cell size. In order of simplest to most complex, these are 1) *the limiting precursor model*, which posits that large cells have higher biosynthetic capacity and, thus, make more organellar building blocks, which are incorporated into more/larger organelles; 2) *the relative growth model*, which posits that cells and organelles follow their own independent growth trajectories; 3) *the demand-driven model*, which posits that organelles grow or multiply to meet the demands of their cell’s size; 4) *the size measurement model*, which posits that the size of the cell is measured, and this measurement is used to set organelle size/number; and 5) *the programmed scaling model*, which posits that cells adjust biosynthesis of organellar building blocks to achieve a match between organelle size/number and target cell size (Marshall 2020). The simplest model that can explain observed patterns in the data is to be favored. Here, we discuss scaling of each organelle and what it suggests about the mechanisms of organellar scaling with cell size.

### Scaling of the Nucleus

Our results on scaling of the nucleus across this ~50-fold increase in cell size are consistent with earlier results in two ways: 1) The nucleus scales allometrically with respect to its shape, showing the decreased SA:V ratio predicted by a sphere increasing in size. 2) The volume of the nucleus scales proportionately with cell size, i.e., the nucleocytoplasmic ratio is maintained, consistent with its apparent functional significance (Balachandra et al. 2022). Mechanistically, the size of the nucleus is set by several interacting forces: DNA content, the state (i.e. compaction) of the chromatin, the total cellular amount of cytoplasmic factors involved in nuclear growth, and overall nucleocytoplasmic transport (Cantwell and Nurse 2019b,a; Heijo et al. 2022; Chen et al. 2023). Across species, the varying contributions of these forces, as well as the differences in the relevant cytoplasmic factors, can be understood in light of a general model for maintenance of the nucleocytoplasmic ratio in which the balance of mechanical forces generated by osmotic pressure within the nucleus and within the cytoplasm sets nuclear volume (Deviri and Safran 2022).

Because of salamanders’ enormous cell volumes, the conserved nucleocytoplasmic ratio yields a disproportionately low nuclear surface area relative to both nuclear and cell volumes. In addition, nuclear pore complexes, which mediate the import and export of macromolecules into and out of the nucleus, are sparse across the salamander nuclear envelope relative to their density in other vertebrates (Maul and Deaven 1977). Nonetheless, the conserved nucleocytoplasmic ratio in salamander cells − as well as overall cell functionality − indicates the existence of effective nucleocytoplasmic transport, demonstrating that movement through the nuclear pore complexes is not a rate-limiting factor for salamander cell physiology, despite proportionally low nuclear surface area and low nuclear pore complex density. Overall, the proportionate increase in nuclear volume in these huge cells is consistent with nuclear scaling following a limiting precursor model.

### Scaling of the Mitochondria

Across the tree of life, across cell sizes that span four orders of magnitude, mitochondrial volume increases proportionately with cell size, comprising roughly 10% of overall cell volume (Okie et al. 2016). Our results are largely consistent with these broad patterns; mitochondrial volume in our representative amphibians ranges from ~9 – 12%. Mitochondrial surface area has also been shown to scale roughly proportionately with cell surface area across a dataset of eukaryotes that includes unicellular organisms, green algae and land plants, and mammals (Lynch and Marinov 2017); our results show somewhat higher mitochondrial surface area per cell surface area than predicted from this broad relationship, but given the sparse sampling and noise inherent in this growing but limited empirical dataset, we do not attach any functional interpretation to this pattern.

In heterotrophic organisms, the increase in overall mitochondrial volume accompanying increased cell volume is primarily accomplished through an increase in mitochondrial number rather than size (Okie et al. 2016). In addition, experimental manipulations in yeast demonstrate that mitochondrial shape stays the same across increasing cell and overall mitochondrial volumes (Seel et al. 2022). Our results showing unchanged mitochondrial SA:V ratio across our model amphibians also suggest isometric growth of the total mitochondrial network with respect to its shape, consistent with these previous results. Overall, our mitochondrial results are broadly consistent with global patterns emerging from limited empirical data across the eukaryotic tree, suggesting that mitochondrial scaling follows general rules that transcend species and cell type.

Despite the presence of general scaling rules suggested by these broad-scale phylogenetic patterns, mitochondrial number is also known to vary at a finer scale among cells, individuals, and species in association with metabolic demand (Schwerzmann et al. 1989; Scott et al. 2018). Salamanders have lower metabolic rates than frogs, and in fact show the lowest metabolic rates among tetrapod vertebrates (Pough 1980; Gatten et al. 1992; Chong and Mueller 2013). Accordingly, they have long been studied with the goal of connecting metabolic rate to cell physiology (Szarski 1970; Goniakowska 1973). The association between low metabolic rate and huge cells in salamanders prompted the “frugal metabolic strategy” hypothesis that natural selection favors large cell size because it relaxes energetic requirements (Szarski 1983) − in contrast to selection at the other physiological extreme favoring small cell size to facilitate high metabolic output in bats, birds, and pterosaurs (Hughes and Hughes 1995; Organ and Shedlock 2009; Wright et al. 2014; Kapusta et al. 2017). Since then, mechanistic hypotheses connecting large cell size to low metabolic rate have been proposed based on both energy supply (e.g. constraints on intracellular resource transport) and demand (e.g. decreased relative cost of Na^+^ – K^+^ gradient maintenance across the plasma membrane) (Glazier 2022).

Despite solid theoretical predictions for lowered metabolic rates in large cells (Kozłowski et al. 2003), empirical data have both supported and rejected a causal relationship between cell size and metabolic rate in amphibians. For example, studies of *Xenopus* frog embryos that differ in cell size and ploidy show lower mass-specific metabolic rates because of decreased relative Na^+^ – K^+^ ATPase activity costs in larger cells (Cadart et al. 2023). In contrast, studies of salamanders fail to show a consistent relationship between mass-specific metabolic rate and cell size’s most commonly-used proxy, genome size (Licht and Lowcock 1991; Gregory 2003; Uyeda et al. 2017; Gardner et al. 2020; Johnson et al. 2021). Our results reveal no effect of lower metabolic rate − whatever its underlying mechanistic cause − on mitochondrial proportionate volume, surface area, or shape in amphibians, suggesting that differences in metabolic demand across species do not drive cellular mitochondrial content in the clade. Our results are more consistent with mitochondrial volume increasing proportionately with cell volume, independent of any specific features of cell physiology.

Mitochondrial content and function can be decoupled (Miettinen and Björklund 2016), and salamander mitochondrial oxidative phosphorylation (OXPHOS) genes show evidence of relaxed purifying selection relative to frogs, consistent with lower functional demand on ATP synthesis (Chong and Mueller 2013). Thus, uniform mitochondrial abundance should not be interpreted as uniform capacity for ATP synthesis across amphibians. More generally, mitochondrial fission and fusion events maintain connectivity among the mitochondria to maximize overall function (Miettinen and Björklund 2016), possibly through the transmission of the proton motive force itself from one region of the mitochondrial network to another faster than the corresponding metabolites and oxygen could reach distant mitochondria by diffusion (Glancy et al. 2015; Miettinen and Björklund 2017). Our results are consistent with mitochondrial content scaling proportionate to cell volume following a limiting precursor model, which yields a mitochondrial network that comprises ~10% of the cell’s volume. This percentage volume, in turn, produces a spatial distribution of organelles that can undergo fission and fusion events to maintain an effective functional network throughout the cell − even at the extremes of animal cell size and metabolic rate.

### Scaling of the Endoplasmic Reticulum and Golgi Apparatus

The largest cell (*N. beyeri*) has significantly less ER surface area density than the smallest cell (*S. tropicalis*). The ER is a complex organelle consisting of different interconnected structures that perform different functions: the nuclear envelope, peripheral cisternae around the nucleus, and a tubular network that extends throughout the cytoplasm (English and Voeltz 2013; Gubas and Dikic 2022). Our analyses were not able to distinguish among these different components of the overall ER, limiting our ability to functionally interpret differences in ER surface area density. However, decreased ER surface area density in the largest cells suggests the possibility that ER functions that take place in the membrane (e.g. membrane lipid synthesis) (English and Voeltz 2013) are operating at lesser capacity per unit of cytoplasm in larger cells relative to smaller cells. This pattern is consistent with the existence of relatively less plasma and nuclear membrane in larger cells (Table 1). In contrast, ER volume scales proportionately with cell volume, suggesting that ER functions that take place in the lumen (e.g. protein folding, processing, and assembly) are operating at the same capacity per unit of cytoplasm in larger and smaller cells. Taken together, these surface area and volume patterns suggest that the overall ER network shape is different in larger cells, trending towards less tubular morphology. Overall, our results are consistent with endoplasmic reticulum content scaling proportionate to cell volume following a limiting precursor model, which yields an ER network that comprises ~6% of the cell’s volume, albeit with decreased SA density in the largest cells that may track lower membrane biosynthesis functional demands. This percentage volume, in turn, produces a spatial distribution of tubules and cisternae that can maintain the physical connections between the ER and the mitochondria, plasma membrane, nucleus, and Golgi apparatus that are necessary for the functionality of all of these organelles (Heald and Cohen-Fix 2014).

The Golgi apparatus receives processed proteins from the ER via membrane-bound vesicles and completes the final stages of the secretory pathway: additional protein processing, sorting, and export in vesicles to the proteins’ final destinations in the cell (e.g. the plasma membrane for secretion or the lysosomes for degradation) (Sengupta and Linstedt 2011). Thus, Golgi size is proximately held at a dynamic equilibrium by balanced influx and outflux of cargo in membrane-bound vesicles, and ER and Golgi size are functionally interconnected (Sengupta and Linstedt 2011; Reber and Goehring 2015). Ultimately, though, Golgi size does scale with cell volume during the growth phase of the mammalian cell cycle (Sin and Harrison 2016), and we show here that Golgi content also scales proportionately with cell volume across evolutionary increases in cell size. Intracellular transport distance is among the most obvious parameters affected by increased cell size; thus, the result that the organelle responsible for transportation out of the cell via the secretory pathway is proportionately the same size across a 50-fold difference in cell volume is noteworthy, suggesting scaling based on a limited precursor model rather than a demand-drive model.

### Organellar Scaling Mechanism Yields Insights into the Proteome of Gigantic Animal Cells

Our overall results are broadly consistent with the limited precursor model; the cytoplasm of larger cells contains more organellar building blocks, which are simply incorporated into larger organelles following the same “rules” across cell sizes, despite differences in metabolic demand and intracellular transport distances. The fact that all organelles scale proportionately with cell volume indicates that all of the proteins required for organelle biosynthesis are maintained at the same concentrations across this evolved ~50-fold difference in cell volume. This finding is significant because it contradicts results from experimentally manipulated increases in cell volume, which reveal proteome dilution and perturbation so severe the cells appear senescent (Neurohr et al. 2019; Cheng et al. 2021; Lanz et al. 2022; Xie et al. 2022). Our results suggest that salamanders have evolved the biosynthetic capacity to maintain a functional proteome despite a huge cell volume. Further work will reveal the dynamics of RNA and protein synthesis and stability that underlie this hypothesized alteration to biosynthetic output.

